# Transertion provides evidence for coupling of transcription and translation in *Bacillus subtilis*

**DOI:** 10.64898/2026.03.21.712414

**Authors:** Jonathan Norris, James Grimshaw, Henrik Strahl, Nikolay Zenkin

## Abstract

In Gram-negative bacteria, co-translational insertion of membrane proteins into the plasma membrane may be coupled to ongoing transcription, a phenomenon known as transertion. Transertion results in a physical shift of the coding gene from the nucleoid towards the membrane and is one of the determinants of the shape of the nucleoid and placement of cell division in Gram-negative bacteria. In contrast, the existence of functional coupling of transcription and translation in *Bacillus subtilis* and potentially other Gram-positive bacteria has been questioned, suggesting that transertion may not happen. Here, by imaging vertically oriented *B. subtilis* cells, we show that the gene of a transmembrane protein changes its localization from inside the nucleoid to the plasma membrane upon induction of its transcription. Localization is restored to the nucleoid when induction of transcription has ceased. The shift of the gene towards the membrane is strictly dependent on transcription, its induction, translation and transmembrane nature of the coded protein. These results suggest that, at least, some principles of cellular regulation based on functional coupling of transcription and translation may be conserved between Gram-positive and Gram-negative bacteria.

## Introduction

In bacteria, transcription and translation can be coupled and occur simultaneously due to the absence of a membrane separating the chromosome from the cytoplasm. The coupled trailing ribosome can functionally control the transcribing RNA polymerase (RNAP) through a direct contact, rescuing it from pausing or terminating stalled RNAPs that are beyond rescue^1–3^. The coupled ribosome can also functionally control transcription termination/antitermination at a distance by affecting the formation of a termination hairpin in the nascent RNA in a phenomenon known as attenuation^4^. Moreover, a premature termination by RNA translocase Rho is prevented by the coupled ribosome that protects transcribing RNAP from Rho^5^.

Coupling of transcription and translation is also a prerequisite for transertion, which functionally links the co-translational insertion of membrane proteins into the plasma membrane to ongoing transcription, thereby physically connecting coding genes to the plasma membrane. Transertion is postulated to be one of the major cellular processes that organize and shape the bacterial nucleoid, keeping it in an expanded state^6–8^. Inhibiting either translation or transcription has pronounced effects on the shape of the nucleoid, which has been argued to result from detaching its connections to the plasma membrane due to disruption of transertion^6^. Furthermore, attachment of the nucleoid to the plasma membrane via coupled RNAP and ribosome has been proposed to determine the placement of the cell division between daughter chromosomes^9–11^.

The above-mentioned examples of functional coupling of transcription and translation were observed only in a number of Gram-negative bacterial species, and transertion was directly observed in *Escherichia coli* and *Vibrio parahaemolyticus*^12–14^. In contrast, transcription and translation in Gram-positive *Bacillus subtilis* were proposed to be functionally uncoupled, because, in some *B. subtilis* genes, transcription was shown to proceed much faster than translation, suggesting that regulation based on coupling is impossible^15^. Furthermore, in *B. subtilis* (and many Gram-positives), transcription termination hairpins of many mRNAs are located very close to the translation stop codons^15^, which means that coupling of transcription and translation may lead to frequent attenuation of transcription termination, which may be harmful to the cell. These observations and considerations suggest that transertion cannot exist in *B. subtilis*, which, in turn, implies that the mechanisms shaping the nucleoid and cell-division placement in *B. subtilis* must differ from those in *E. coli*.

The main pathway for co-translational incorporation of transmembrane proteins into the plasma membrane in *B. subtilis* (and bacteria in general) is via the SecYEG translocase^16,17^. Translation of the transmembrane protein is paused by Signal Recognition Particle (SRP), which recognizes a hydrophobic signal anchor emerging from the ribosome. The paused translation elongation complex is then transferred to the SecYEG translocase at the membrane. In this study, we investigated whether the incorporation of transmembrane proteins into the plasma membrane involves transertion in *B. subtilis*, that is, whether ribosomes synthesizing transmembrane proteins are coupled to transcription, leading the corresponding gene to shift towards the membrane (Fig. 1A). To investigate transertion in *B. subtilis*, we chose to focus on the cold shock response operon containing three genes: *des, desK* and *desR* (Fig. 1A, B). The fatty acid desaturase, Des, is a transmembrane protein that is expressed in response to cold shock conditions to increase fatty acid unsaturation and, thus, to re-fluidize the membrane^18,19^, and that is inserted into the plasma membrane in a co-translational manner via SRP/SecYEG pathway^20,21^. During cold shock, the sensory kinase DesK phosphorylates DesR, which activates transcription from the P*des* promoter and the corresponding synthesis of Des. Upon restoration of membrane fluidity, DesR is dephosphorylated by DesK, leading to stop of transcription from P*des*^*18*^.

**Figure 1.**
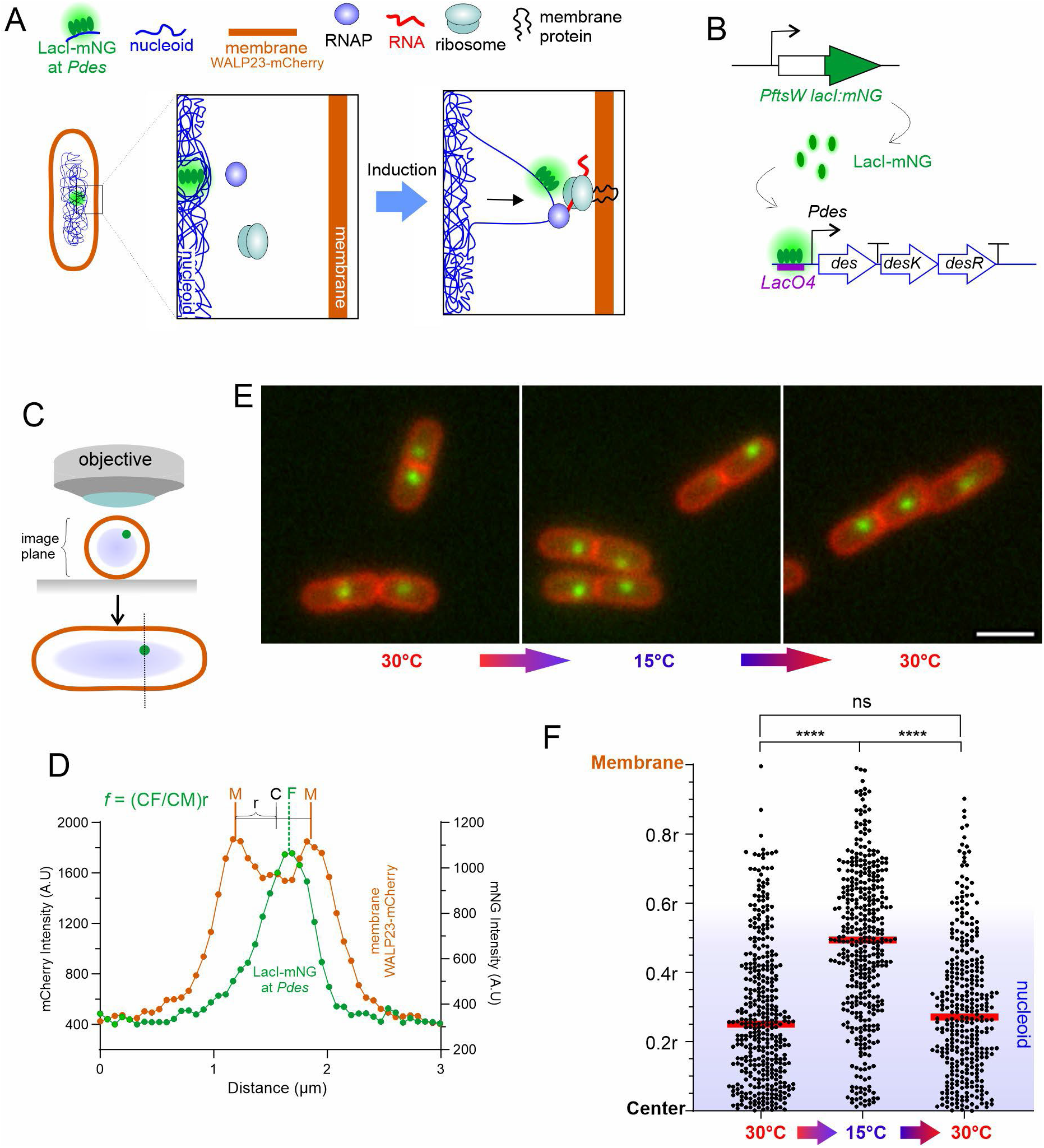
Visualization of transertion. **A**. Cartoon depicting the principle of transertion, yet hypothetical in *B. subtilis*. LacI-mNG and WALP23-mCherry were used to visualize genomic locus and the membrane, respectively. **B**. Scheme of fluorescent labelling of the *des* operon upstream of the *des* promoter (P*des*) by *lacO*-LacI-mNG. **C and D**. Quantitation of focus position during imaging of horizontal cells. The position of *des* focus was determined as a fraction of cell radius (r) from the cell center towards the cell membrane. **E**. Horizontally imaged cells grown at 30°C, cold shocked for 15 min, and recovered from cold shock at 30°C for 30 min. Green focus is *des* (labelled with *lacO*-LacI-mNG); red is membrane labelled with WALP23-mCherry (see panel A). **F**. Quantification of *des* foci distribution before cold shock (n=387), after cold shock at 15°C (n=401) and following recovery at 30°C (n=343). The red line represents the median value. The approximate position (due to irregular shape) of the nucleoid was determined with Hbs-GFP.

## Results

### Movement of genomic DNA locus towards membrane

To visualize potential transertion of *des*-locus, we fluorescently labelled chromosomal DNA upstream of *Pdes* (85 base pairs upstream of transcription start site) by introducing an array of four Lac-repressor binding sites (*LacO4*) and expressing Lac repressor fused to fluorescent mNeonGreen (LacI-mNG) *in trans* (Fig. 1A, B). The position of the fluorescent focus upstream of *des* gene (usually one per cell given the terminus-proximal location of *des*-locus) was then visualized using fluorescence microscopy (Fig. 1C, D). The plasma membrane was visualized in the same cells by expressing a model transmembrane domain, WALP23, fused to mCherry (WALP23-mCherry; Fig. 1A). Position of the fluorescent focus was calculated as a fraction of cell radius (r) from the center of the cell towards the membrane (Fig. 1C-E).

At 30°C, the median of distribution of *des* foci was at 0.25r (Fig. 1F). Cooling cells from 30°C to 15°C (cold shock for *B. subtilis*) for 15 min resulted in a significant shift of the foci towards the membrane to 0.49r (Fig. 1F). Moreover, recovering cells from cold shock by returning them to 30°C for 30 min resulted in a return of the foci to the original position (median at 0.27r; Fig. 1F).

Imaging horizontal cells biases the observed distribution of the foci towards the cell center because the position of a focus along the axis of observation is projected onto a 2D plane, causing peripheral signals to appear more central than they are in a 3D space. We therefore applied imaging of vertically oriented cells – VerCINI^22^ (Fig. 2A). Here the cells are placed in vertical nanoholes, with imaging taking place along the long axis of the cells (Fig. 2A). Now, the observed position of the foci (unless the focus moves towards a pole of the cell) reflects the actual distances from cell center towards the membrane (Fig. 2A). As above, the positions of a *des* foci were calculated as a fraction of cell radius (r) from the center of the cell towards the membrane (Fig. 2B, C).

**Figure 2.**
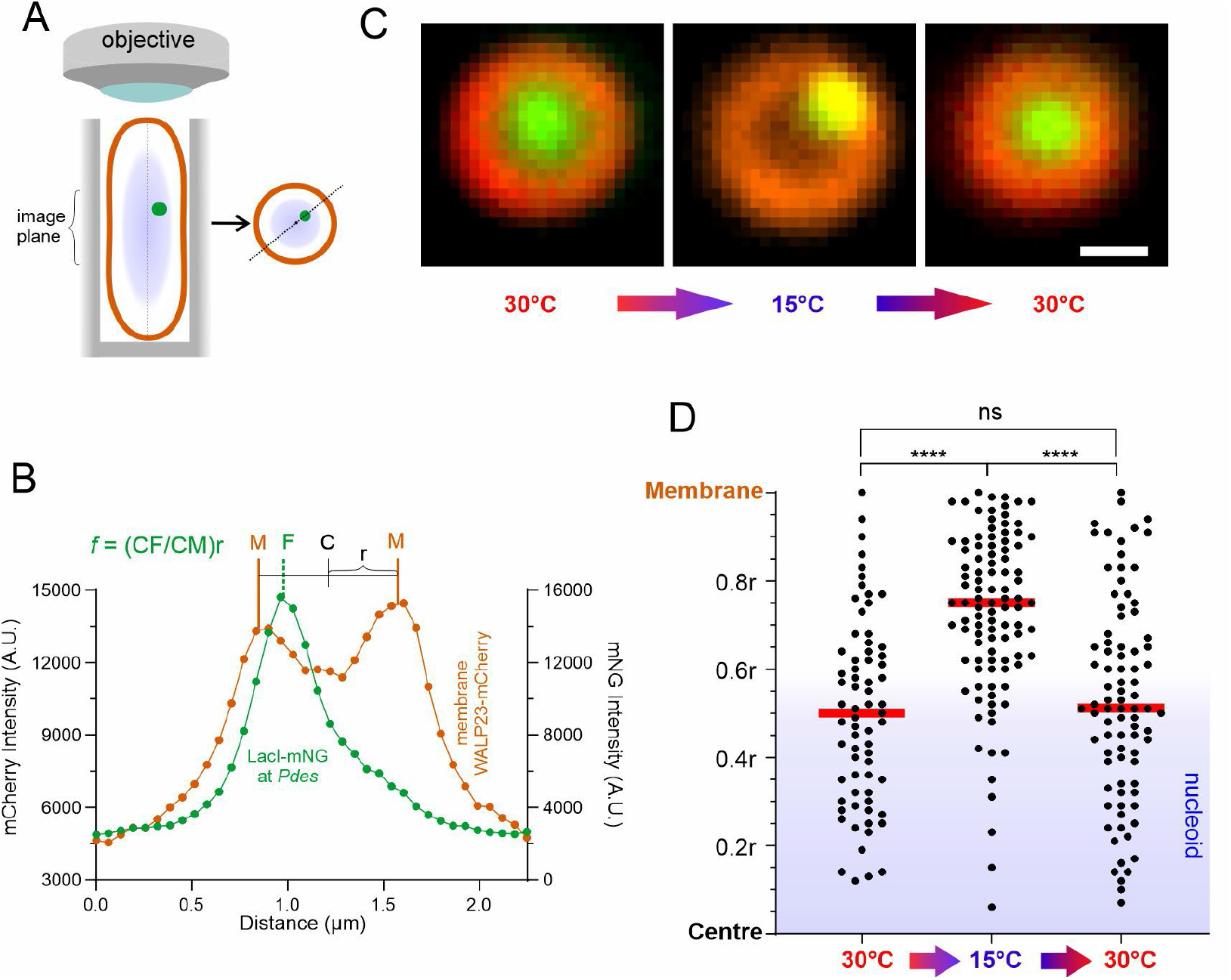
Visualization of transertion using vertically imaged cells. **A**. VerCINI, a technique of imaging vertically oriented bacterial cells along their long axis, eliminates the bias of fluorescent foci positioning relative to the center of the cell and the membrane. Cells are positioned vertically inside nanoholes in the agarose pad (grey). **A and B**. The position of *des* focus is determined as a fraction of the cell radius (r) from the cell center towards the cell membrane. **C**. Representative images of vertically imaged cells grown at 30°C, cold shocked for 15 min, and recovered from cold shock at 30°C for 30 min. Green focus is *des* (labelled with *lacO*-LacI-mNG); red is membrane labelled with WALP23-mCherry (see Fig. 1A). **D**. Quantification of *des* foci distribution before cold shock (n=69), after cold shock at 15°C (n=100) and following recovery at 30°C (n=86). The red line represents the median value.

As can be seen from Fig. 2D, at 30°C, the median of *des* foci positions was at 0.5r. Cold shock at 15°C resulted in a shift of the median towards the membrane to 0.74r (Fig. 2D). Recovering cells from the cold shock resulted in *des* returning to its original position (0.5r; Fig. 2D).

### DNA locus shift depends on the expression of the gene and on the transmembrane nature of the protein product

Deletion of the binding site for the *des* activator DesR prevented movement of *des* foci towards the membrane upon the cold shock (Fig. 3A), strongly arguing that the observed shift depends on transcriptional activation. Moreover, wild-type cells (with respect to regulation of P*des*) adapted to 15°C for 72 hours and, thus, no longer expressing *des*^23,24^, featured *des* foci at the same position as in the cells grown at 30°C (Fig. 3B).

**Figure 3.**
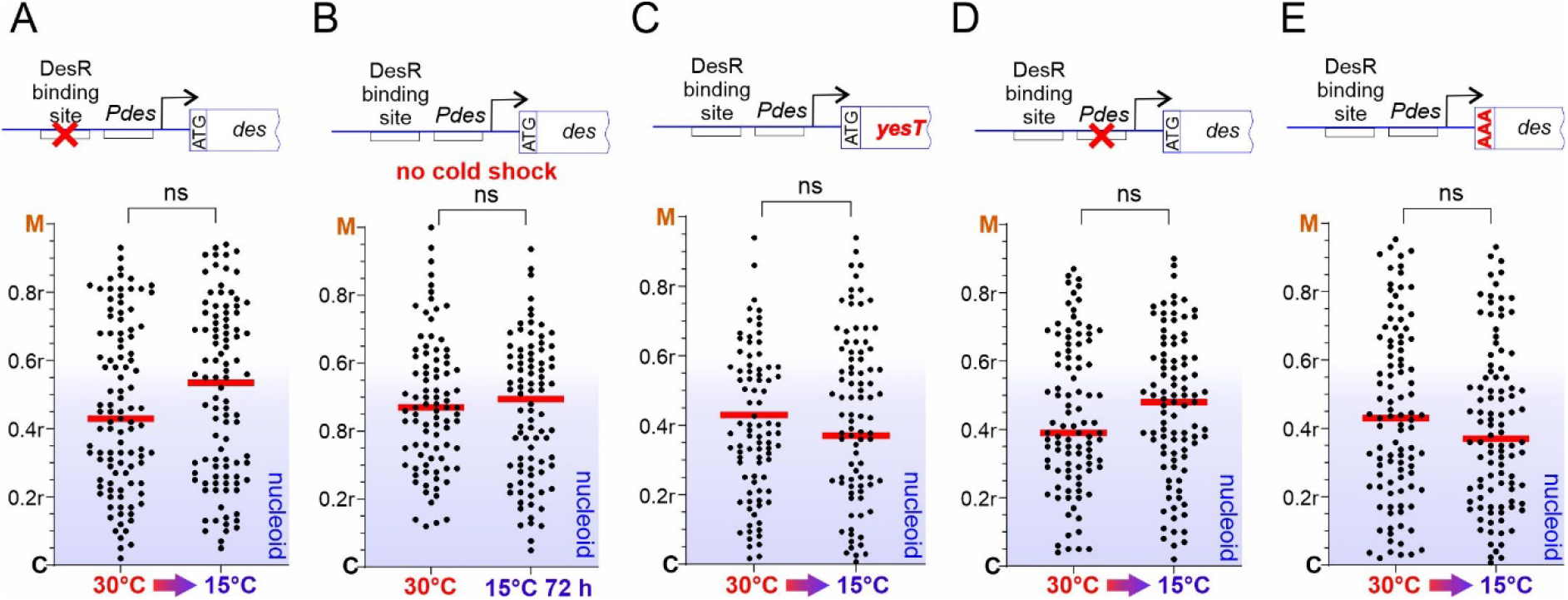
Genetic and condition determinants of transertion. *des* foci distribution in: **A**. cells with DesR binding site removed (n=62 at 30°C; n=97 after cold shock); **B**. cells after continuous growth at 30°C (n=89) or 15°C (n=87) (no cold shock). **C**. cells with *des* gene substituted for a gene coding for cytoplasmic protein YesT (n=87 at 30°C; n=84 after cold shock); **D**. cells with *des* promoter (P*des*) removed (n=97 at 30°C; n=96 after cold shock), **E**. cells with *des* start codon mutated (n=98 at 30°C; n=100 after cold shock).

Transertion depends on the SRP/SecYGF mechanism that facilitates transmembrane protein insertion into the membrane^16,17^. To test whether the observed movement of *des* foci towards the membrane depends on the transmembrane nature of Des, we replaced the coding sequence of the *des* gene with that of a cytoplasmic protein *yesT*. As seen from Fig. 3C, the cold shock did not affect the position of *yesT* expressed from the *des*-locus.

To directly show if *des* foci movement depends on transcription and translation of *des* gene, we mutated either *Pdes* promoter (Fig. 3D) or the start codon of *des* open reading frame (Fig. 3E). Either of the mutations prevented movement of the *des* foci during cold shock (Fig. 3D,E). Importantly, the above results also confirm that the temperature shift alone is not responsible for the movement of genomic DNA towards the membrane.

## Discussion

Transertion, i.e. coupling of transcription to translation to protein insertion into the membrane, has been so far only observed only in Gram-negative bacteria^12–14^. Moreover, the proposed functional uncoupling of transcription and translation in Gram-positive bacteria^15^ implies that transertion may not happen in Gram-positive bacteria. The principal finding of our work is the demonstration of the transertion phenomenon in the Gram-positive model organism *B. subtilis*. This suggests that transertion may be conserved among all bacteria, which has the following important implications:

1. Our observation of transertion in *B. subtilis* indicates that the speeds of RNAP and ribosome may be coordinated, at least on some genes, and, thus, transcription and translation can be functionally coupled like in Gram-negative bacteria. This, in turn, suggests that mechanisms of mutual regulation between coupled transcription and translation that are common for Gram-negative bacteria^2,25^ may also exist in Gram-positives. These observations challenge the previous proposal of global functional uncoupling of transcription and translation in *B. subtilis*, and suggest that both coupled and uncoupled modes may coexist depending on the genetic context. It is possible that the coupling of transcription and translation depends on the nature of the protein product. Des is a transmembrane protein and requires special arrangements, such as transertion, during its synthesis, while previously studied genes code for non-transmembrane proteins (*tkt* and *pyrA*)^15^.
2. The expanded shape of nucleoid in Gram-negative bacteria is thought to be mainly determined by transertion at many genes coding for transmembrane proteins, which “pulls” on the nucleoid towards the cell periphery^8,26^. Our results indicate that the expanded shape of the nucleoid of Gram-positive *B. subtilis*, and possibly other Gram-positives, may also be driven by transertion.
3. In *B. subtilis*, the placement of the division septum is determined by the position of the daughter chromosomes, which interact with the membrane through protein Noc that binds both DNA and membrane^11,27^, and the Min-system that inhibits cell division at the cell poles and close to the active septum^28^. Because the deletion of *noc* did not fully disrupt the septum placement^29^, it was hypothesized that transertion may act synergistically with Noc in determining the placement of the division septum^11^. Observation of transertion in *B. subtilis* supports this hypothesis.

How abundant transertion and functional coupling between transcription and translation are in *B. subtilis* and other Gram-positive species is yet to be understood. However, our findings suggest that general mechanisms underlying the functional coupling of transcription and translation may be conserved across Gram-positive and Gram-negative bacteria, although the prevalence of coupled and uncoupled modes may differ between the phyla.

## Methods

### Plasmid construction

Plasmids were constructed using standard molecular cloning techniques and confirmed by sequencing. The plasmids employed in this study are listed in Table S1, while the primers used for cloning are listed in Table S2.

pJN11, which constitutively expresses lacI-mNeonGreen, was constructed by inserting *amyE::PftsW lacI-mNeonGreen spec* synthetic construct (IDT) into pUC18 linearized by oligos JN13 and JN14. This plasmid and all further plasmids were constructed using NEBuilder, utilizing NEBuilder assembly tool and methodology. pJN26, which contains the *des* promoter flanked by 1kb of sequence on either side, was generated by amplifying the insert (from *B. subtilis* 168 chromosomal DNA with oligos JN17 and JN16) and inserting it into pUC18 linearized via oligos JN52 and JN53. pJN27, encoding 4 repeating *lac* operator sequences with a kanamycin cassette 85bp upstream of the *des* promoter, was generated by inserting *lacO4* (amplified from *B. subtilis* strain BWX1200^30^ with oligos JN28 and JN29) and kanamycin resistance cassette (amplified from *B. subtilis* strain BWX1200 with oligos JN23 and JN24) into pJN26 linearized with oligos JN38 and JN39. pJN207, containing the *lacO4* upstream of *des* promoter with mutated *des* start codon (ATG→AAA), was generated via site-directed mutagenesis on plasmid pJN27 with oligos JN133 and JN134. pJN210, containing the *lacO4* upstream of the deleted *des* promoter site, was generated using plasmid pJN27 via site-directed mutagenesis with oligos JN149 and JN150. pJN212, containing the *lacO4* upstream of *des* promoter with a deletion of the DesR binding site sequence, was generated using plasmid pJN27 via site-directed mutagenesis with oligos JN153 and JN154. pJN220, containing the *lacO4* upstream of *des* promoter with *des* replaced by *yesT*, was generated by inserting *yesT* gene (amplified with primers JN139 and JN140 from *B. subtilis* 168 genomic DNA) into plasmid pJN27 (linearized using primers JN137 and JN138).

### Strain and plasmid construction

Bacterial strains used in this study and details of their construction are listed in Table S3. All genetic constructs were verified by sequencing.

### Fluorescence microscopy of horizontal cells

Overnight cultures were inoculated and incubated at 30°C shaking at 175 RPM in LB media supplemented with 0.2% glucose. Next day, overnight cultures were 1/100 diluted into fresh SMM microscopy media (Table S4). Cultures were grown until the desired optical density, which for most experiments was OD_600_ of 0.3. Unless stated otherwise, cold shock was performed on culture samples mounted on microscopy slides prepared in SMM medium, and assembled as detailed in de Jong et al ^31^. For this aim, the slides were incubated at 15°C for 15 min, followed by rapid imaging at room temperature before the sample had time to warm up. Fluorescence microscopy was carried out with Nikon Ti2 using NIS Element imaging software, CoolLed pE-4000 light source, Photometrics Kinetix camera, and Nikon CFI Plan Apo Lambda 100x/1.45 NA Oil Ph3 objective. mNeonGreen constructs were imaged using Semrock GFP-4050B filter cube (EX466/40, DM495lp, EM525/50) and 470 nm LED, and mCherry constructs using a Semrock mCherry-C (EX562/40, DM593lp, EM641/75) and 580 nm LED. Where applicable, the microscope chamber was set to the experimental temperature. Since 15°C was not achievable, the microscope chamber was, in this case, left at room temperature.

### Vertical cell imaging by nanostructured immobilization (VerCINI)

Overnight cultures were prepared as described above. The next day, 1/100 dilutions from overnight cultures were transferred into fresh SMM microscopy media (Table S4). Cultures were grown at 30°C (except for the experiment shown in Fig. 3B, in which cells were grown at 15°C) until OD_600_=0.3. For vertical immobilization, 800 µl of 6% solution of ultrapure agarose in SMM microscopy media was applied onto a custom micro-pillar wafer^22^. A glass slide with a GeneFrame (Thermo Fisher Scientific) mounted on it was pushed down onto the wafer for 20 seconds until the agarose polymerized. 1 ml of cells at OD_600_ = 0.3 were transferred into 1.5 ml Eppendorf test tubes, then spun down for 1 min at 13.000 x *g*, supernatant removed, and cell pellet suspended in SMM microscopy media. 8 µl of cells were added to the imprinted agarose. These steps were completed at either 30°C or 15°C, respectively. Slides were then centrifuged on a flat-bottomed 96-well plate rotor for 4 minutes at 4000 x *g*. Non-trapped cells were washed away with the same media. Samples were left to air-dry for 3 minutes before removing non-imprinted agarose regions to create air pockets providing O_2_-supply, followed by sealing with a coverslip.

VerCINI^22^ was carried out with Nikon TI2 equipped with Nikon CFI SR HP Apo TIRF 100x/1.49 NA Oil objective, 488nm and 561nm solid state laser sources for the excitation of mNeonGreen and mCherry, respectively, and imaged using Photometrics Kinetix camera and NIS Elements imaging software. The imaging filter sets used were Chroma TRF49904 (EX488/10, DM488lp, EM525/50 + 500lp) Chroma TRF49909 (EX561/10, DM561lp, EM600/50 + 575lp). Highly inclined and laminated optical sheet microscopy (HiLO) imaging, which increases the signal-to-background noise ratio by inclining the laser beam to penetrate the specimen at an angle, was used. As a result, illumination of the specimen is laminated as a thin optical sheet^32^.

### Horizontal Image analysis

Image’s taken for horizontal cells were analyzed and quantified utilizing programs Ilastik^33^ and FIJI^34^ Images were corrected for chromatic aberration between wavelengths of red and green channels, with a scaling factor of 1.0018 applied to red channel. Cells were segmented via Ilastik using the membrane channel. To locate *lacO4*/LacI foci in images, Trackmate’s Laplacian of Gaussian (LoG) detector was used ^35,36^ with an estimated foci diameter of 0.2 μm and a minimum quality threshold of 500.

Foci positioning was determined within the context of the cell by bisecting the focus with lines passing through the membrane, iterated at 10-degree intervals. A fluorescence intensity profile was measured for the membrane channel, and then a Gaussian fit was applied. Mid-cell was determined by the centroid and membrane edges from the full width half maximum (FWHM) of the Gaussian curve. Foci distance was then calculated relative to these values.

### Vertical Image analysis

Image analysis was performed in FIJI ^34^ Images were corrected for chromatic aberration between the red and green channels, with a scaling factor of 1.0018 applied to the red channel. Cells were segmented via local thresholding in FIJI via Bersen local thresholding algorithm ^37,38^ creating a binary image. Individual cell ROIs were generated from the resulting binary image with the Analyze Particles function in Fiji, filtering for a circularity between 0.8 and 1.0 and an area of greater than 0.5 μm^2^. To locate *lacO4*/LacI foci in images, Trackmate’s Laplacian of Gaussian (LoG) detector was used ^35,36^ with an estimated foci diameter of 0.2 μm and a minimum quality threshold of 500.

To calculate the distance of a focus from the membrane, an intensity profile of a line running from the centroid of the cell ROI through the focus coordinates and across the membrane of the cell was measured. Membrane position was then determined by fitting a Gaussian curve, with the membrane being identified as the peak of the Gaussian. The distance from the centroid to the membrane (cell radius) was doubled to output cell width. Output of foci distances towards the membrane was calculated as a fraction of the radius. In addition, the data has been manually validated.

Analysis of strain BJN047 was achieved manually in FIJI. Images were corrected for chromatic aberration between red and green channels, scaled by a factor of 1.0018 to the red channel. Cells were selected and manually bisected by a straight line. A fluorescence intensity line scan of the membrane and nucleoid stains revealed an approximate average edge of the nucleoid, shown as a blue section in plots.

### Statistical analysis

Statistics were determined on graphs using GraphPad Prism 10 via an unpaired t-test using nonparametric Mann-Whitney test.

### Source Code availability

Code mentioned above for horizontal and vertical cell foci analysis is available at https://github.com/NCL-ImageAnalysis/FociToMembraneDistance.git

## Acknowledgements

This work was supported by Wellcome Trust Investigator Award [grant number 217189/Z/19/Z] to NZ and BBSRC equipment Grant [grant number BB/T017570/1] for NZ and HS. George Merces is acknowledged for helping with the quantification of preliminary data that led to this project.

## Data and materials availability

Source data for all figures and graphs presented in the manuscript will be made available via Newcastle University’s research data repository (submission pending). The strains and plasmids are available upon request to NZ and HS.

