## Supplementary Tables for "Transertion provides evidence for coupling of transcription and translation in *Bacillus subtilis*"

**Membrane localization of gene coding for transmembrane protein provides evidence for transertion in *Bacillus subtilis***

**Authors:** Jonathan Norris, James Grimshaw, Henrik Strahl* and Nikolay Zenkin*

**Table S1: List of plasmids**

**Table S2: List of oligonucleotides**

**Table S3: List of bacterial strains**

**Table 4: List of medias and buffers used with compositions.**

**Table S1: List of plasmids.**

| **Plasmid** | **Genotype** | **Source or construction** |
|---|---|---|
| pUC18 | *ori: pMB1 bla lacZ* | (Taylor et al., 1993) |
| pJN11 | *bla amyE::PftsW lacI-mNeonGreen spec* | This work |
| pJN26 | *bla* P*des des* | This work |
| pJN27 | *bla ΩlacO4* P*des des kan* | This work |
| pJN207 | *bla ΩlacO4* P*des* $des^{AUG \rightarrow AAA}$ *kan* | This work |
| pJN210 | *bla ΩlacO4 Δ*P*des des kan* | This work |
| pJN212 | *bla ΩlacO4* P$des^{\Delta DesRbinding}$ *des kan* | This work |
| pJN220 | *bla ΩlacO4* P*des des::yesT kan* | This work |

**Table S2: List of oligonucleotides**

| Name | Sequence |
|---|---|
| JN13 | TGATCCGCTGCCTGATC |
| JN14 | ATGTCCAGACTTCAGATCCACTAG |
| JN16 | TAGATGTCTGCTG+AGCTGCATAGCCGATTGCGCTAAAAAAG |
| JN17 | CGTGATGTTCATGAGCAACTTCAAGC |
| JN23 | CTAAAACAATTCATCCAGTAAAATATAATATTTTATTTTC |
| JN24 | CAATTACCCGTACATAATTGAATCGGCTCCGTAGATAC |
| JN28 | CAATTATGGACGGGTAATTG |
| JN29 | GAATTCGGATCCCTCGAG |
| JN33 | GCTGAATCGAGATCCGAGTGTGC |
| JN34 | CAATAAAGCGTCCTCTTGTG |
| JN38 | GCTCATGAACATCACGCCGAGCTCGAATTCGTAATCATGG |
| JN39 | CCGGATTTCGTCTCCCTGGCCGTCGTTTTACAACG |
| JN133 | GAGAGGATACTTACTAAAACTGAACAAACCATTGC |
| JN134 | GCAATGGTTTGTTCAGTTTTAGTAAGTATCCTCTC |
| JN137 | AGTAAGTATCCTCTCATTGTGTGTC |
| JN138 | TGATAAAGGAGGTCTTCTAATATACTAGAAGG |
| JN139 | GTGATGAAGAAACCGATTCAAGTATTTTTAGC |
| JN140 | TTATACATGTTCTTTTCCCTCCCGGGCAACTTCAAG |
| JN149 | GATGTGTGCTACTACAAAAGACGAACCG |
| JN150 | CCTCTCATTGTGTGTCTCGGTTCGTCTTT |
| JN153 | CGCACGAGACATGACAAATGTGTGTG |
| JN154 | GTCTTTTGTAGTAGCACACACATTTG |

**Table S3: List of bacterial strains.**

Transformation of plasmids into a background strain or crossover events are represented as X>Y.

| Strain | Species | Genotype | Source or construction |
|---|---|---|---|
| 168ca | *B. subtilis* | *trpC2* | (Barbe et al., 2009) |
| BWX1200 | *B. subtilis* | *spoIIIE36*, *yycR*(−7°)::tetO48 cat, *pelB*(+174°)::lacO48 kan, *ycgO*::PftsW *tetR*-cfp spec terminators *PftsW lacI-mypet* | (Wang & Rudner, 2014) |
| BWX721 | *B. subtilis* | *yycR*(−7°)::tetO48 erm, *ycgO*::*PftsW tetR-cfp* spec, *sacA*::*hbs-mypet* kan | (Wang & Rudner, 2014) |
| PL070 | *B. subtilis* | *AprE::Pxyl*$_{constituative}$*WALP23-mCherry cat* | Lee, unpublished |
| BJN004 | *B. subtilis* | *trpC2 amyE::PftsW lacI-mNeonGreen spec* | pJN11>168ca. |
| BJN007 | *B. subtilis* | *trpC2 amyE::PftsW lacI-mNeonGreen spec, AprE::Pyxl WALP23-mCherry cat* | BJN004>PL070 |
| BJN015 | *B. subtilis* | *trpC2 ΩlacO4* Pdes *des kan* | pJN27>168ca |
| BJN016 | *B. subtilis* | *trpC2 ΩlacO4* Pdes *des kan, amyE::PftsW lacI-mNeonGreen spec* | pJN27>BJN004 |
| BJN017 | *B. subtilis* | *trpC2 ΩlacO4* Pdes *des kan, amyE::PftsW lacI-mNeonGreen spec, AmyE::Pyxl WALP23-mCherry cat* | pJN27>BJN007 |

| | | | |
|---|---|---|---|
| BJN047 | *B. subtilis* | *trpC2 sacA::hbs-GFP kan, AmyE::Pyxl WALP23-mCherry cat* | PL070>BWX721 |
| BJN055 | *B. subtilis* | *trpC2 ΩlacO4* P*des* $des^{AUG \rightarrow AAA}$ *kan, amyE::PftsW lacI-mNeonGreen spec, AmyE::Pyxl WALP23-mCherry cat* | pJN207> BJN007 |
| BJN057 | *B. subtilis* | *trpC2 ΩlacO4 Δ*P*des des kan, amyE::PftsW lacI-mNeonGreen spec, AmyE::Pyxl WALP23-mCherry(cat* | pJN210> BJN007 |
| BJN060 | *B. subtilis* | *trpC2 ΩlacO4* P*des des::yesT kan, amyE::PftsW lacI-mNeonGreen spec, AmyE::Pyxl WALP23-mCherry cat* | pJN220> BJN007 |
| BJN062 | *B. subtilis* | *trpC2 ΩlacO4* P$des^{\Delta \mathrm{DesRbinding}}$ *des kan, amyE::PftsW lacI-mNeonGreen spec, AmyE::Pyxl WALP23-mCherry cat* | pJN212> BJN007 |
| DH5α | *E. coli* | *fhuA2 Δ(argF-lacZ)U169 phoA glnV44 Φ80 Δ(lacZ)M15 gyrA96 recA1 relA1 endA1 thi-1 hsdR17* | (Taylor et al., 1993) |

**Table 4: List of medias and buffers used with compositions.**

| Media | Components |
|---|---|
| LB | 10 g Tryptone<br>5 g yeast extract<br>10 g NaCl<br>1 L Distilled water |
| SMM basic salts | 0.2% $(NH_4)_2SO_4$<br>1.4% $K_2HPO_4$<br>0.6% $KH_2PO_4$<br>0.2% $C_6H_5Na_3O_7.2H_2O$ |
| SMM transformation media | 10 ml SMM basic salts<br>120 µl 40% glucose solution<br>60 µl 1M MgSO4<br>100 µl of 2% tryptophan<br>10 µl 20% Casamino acids<br>10 µl of 2.2mg/ml Ferric-ammonium citrate solution. |
| SMM Dilution Media | 10 ml SMM basic salts<br>120 µl of 40% glucose<br>60 µl 1M MgSO4 |
| SMM microscopy media | 10 ml SMM basic salts<br>240 µl 40% glucose solution<br>60 µl 1M MgSO4<br>100 µl of 2% tryptophan<br>10 µl 20% Casamino acids<br>100 µl of 2.2mg/ml Ferric-ammonium citrate solution. |
| TBE (5x) | 54 g Tris base<br>27.5 g boric acid<br>20 ml 0.5M EDTA (pH 8.0) |

Barbe, V., Cruveiller, S., Kunst, F., Lenoble, P., Meurice, G., Sekowska, A., Vallenet, D., Wang, T., Moszer, I., Médigue, C., & Danchin, A. (2009). From a consortium sequence to a unified sequence: the Bacillus subtilis 168 reference genome a decade later. *Microbiology*, *155*(6), 1758-1775. https://doi.org/10.1099/mic.0.027839-0

Taylor, R. G., Walker, D. C., & McLnnes, R. R. (1993). E.coli host strains significantly affect the quality of small scale plasmid DNA preparations used for sequencing. *Nucleic Acids Research*, *21*(7), 1677-1678. https://doi.org/10.1093/nar/21.7.1677

Wang, X., & Rudner, D. Z. (2014). Spatial organization of bacterial chromosomes. *Current Opinion in Microbiology*, *22*, 66-72. https://doi.org/10.1016/j.mib.2014.09.016
